# An ancient fecundability-associated polymorphism creates a new GATA2 binding site in a distal enhancer of *HLA-F*

**DOI:** 10.1101/245043

**Authors:** Katelyn M. Mika, Xilong Li, Francesco J. DeMayo, Vincent J. Lynch

**Affiliations:** Department of Human Genetics, The University of Chicago, 920 E. 58th Street, CLSC 319C, Chicago, IL 60637, USA.; National Institute of Environmental Health Sciences, Department Reproductive & Developmental Biology Laboratory,111 TW Alexander Drive, Research Triangle Park, NC, 27709, USA

## Abstract

Variation in female reproductive traits such as fertility, fecundity, and fecundability are heritable in humans, but identifying and functionally characterizing genetic variants associated with these traits has been challenging. Here we explore the functional significance and evolutionary history of a G/A polymorphism of SNP rs2523393, which we have previously shown is an eQTL for the *HLA-F* gene and significantly associated with fecundability (time to pregnancy). We replicated the association between rs2523393 genotype and *HLA-F* expression using GTEx data and demonstrate that *HLA-F* is up-regulated in the endometrium during the window of implantation and by progesterone in decidual stromal cells. Next, we show that the rs2523393 A allele creates a new GATA2 binding site in a progesterone responsive distal enhancer that loops to the *HLA-F* promoter. Remarkably, we found that the A allele is derived in the human lineage, that G/A polymorphism arose before the divergence of modern and archaic humans, and is segregating at intermediate to high frequencies across human populations. Remarkably, the derived A is also has been identified in a GWAS as a risk allele for multiple sclerosis. These data suggests that the polymorphism is maintained by antagonistic pleiotropy and a reproduction-health tradeoff in human evolution.

## Introduction

Female reproductive traits such as fertility, fecundity, and fecundability are heritable in humans ^1,2^ however, identifying the genetic bases for these traits has been challenging ^1–3^. We previously performed an integrated expression quantitative trait locus (eQTL) mapping and association study to identify eQTLs in mid-secretory endometrium that influence pregnancy outcomes ^1,2,4–6^. Among the eQTLs we identified was a G/A polymorphism (rs2523393) that was significantly associated with *HLA-F* expression and fecundability. Specifically, the G allele of rs2523393 was associated with longer intervals to pregnancy and lower expression of the *HLA-F* in mid-secretory phase (receptive) endometrium, whereas the A allele was associated with shorter intervals to pregnancy and higher *HLA-F* expression. The median time to pregnancy, for example, was 2.3, 2.6, and 4.9 months among women with the AA, GA, and GG genotypes, respectively ^1–3,6^.

While the functions of HLA-F are enigmatic, it is thought to regulate immune responses and may play an important role in regulating maternal-fetal immunotolerance during pregnancy^7,8^. HLA-F is highly expressed, for example, in placental villi, on the surface of invasive and migratory extra-villus trophoblasts (EVTs), and decidual stromal cells (DSCs) ^9–11^. Furthermore HLA-F expression increases during gestation, reaching a peak at term^9,12^. Like other MHC-I molecules, HLA-F binds natural killer (NK) cell receptors from the LIR and KIR families, including LIR1 and LIR2^13,14^, KIR2DS4 and KIR3DL2 ^15^, and KIR3DL1 and KIR3DS1^14,16,17^. Uterine natural killer (uNK) cells, which are essential for the establishment and maintenance of maternal immunotolerance and spiral artery remodeling, express KIR3DL1 and LIR2 suggesting HLA-F expressed by EVTs and DSCs mediate interactions with uNK during implantation, trophoblast invasion, and establishment of the uteroplacental circulation. HLA-F expression level is also positively correlated with uNK abundance in mid-luteal endometria and is predictive of achieving pregnancy^6,18^ consistent an important role for HLA-F in the establishment of pregnancy.

Here we explore the functional and evolutionary history of the rs2523393 G/A polymorphism. We first replicate the association between rs2523393 genotype and *HLA-F* expression using GTEx data. We demonstrate that *HLA-F* expression increases during the menstrual cycle and is up-regulated by progesterone during the differentiation of endometrial stromal fibroblasts into decidual stromal cells. Next, we show that rs2523393 is located within a cAMP/progesterone responsive enhancer that makes long range regulatory interactions with the promoter of *HLA-F*, and that the A allele creates a new binding site for the progesterone receptor (PGR) co-factor GATA2. Remarkably the G/A polymorphism arose before the divergence of modern and archaic humans, is segregating at intermediate to high frequencies across human populations, and associated with a predisposition to several diseases. These data suggest the G/A polymorphism is maintained by antagonistic pleiotropy and that there is a reproduction-health tradeoff in human evolution.

## Materials and Methods

### rs2523393 is an eQTL for HLA-F

We replicated the association between the G/A polymorphism at rs2523393 and *HLA-F* expression levels using GTEx Analysis Release V6 (dbGaP Accession phs000424.v7.p2) data for 35 tissues, including the 101 uterus samples^19,20^. We also used GTEx data to identify other genes for which rs2523393 was an eQTL. Briefly, we queried the GTEx database using the ‘Single tissue eQTLs search form’ for SNP rs2523393.

### *HLA-F* is expressed in decidual stromal cells and is differentially regulated by progesterone

Using previously generated microarray expression data, we examined the expression of HLA-F in the endometrium across the menstrual cycle (GSE4888^21^) and in endometrial samples from fertile women (n=5), woman had implantation failure following IVF (n=5), and woman with recurrent spontaneous abortion (n=5) (GSE26787^22^). These microarray datasets were analyzed with the GEO2R analysis package, which implements the GEOquery ^23^ and limma R packages ^24,25^ from the Bioconductor project to quantify differential gene expression. We also examined *HLA-F* expression in RNA-Seq data previously generated from ESFs treated with control media or differentiated (decidualized) with 100nM medroxyprogesterone acetate, and 1mM 8-bromo-cAMP (all from Sigma-Aldrich Co., St. Louis, MO) into DSCs ^26,27^. We also used previously generated RNA-Seq data to characterize the expression level of *HLA-F* in human endometrial stromal fibroblasts (ESF), decidual stromal cells (DSCs), and DSCs treated with an siRNA to knockdown PGR expression ^38^.

### GATA2 ChIP-Seq

ESFs were cultured in 150-mm culture dishes with HESC growth media (DMEM/F12 containing 10% of fetal bovine serum and 1% antibiotic-antimycotic). Cells were grown to each approximately 90% confluency, starting to treat with decidual media (Opti-MEM media containing 2% charcoal-stripped FBS, 1% antibiotic-antimycotic, 1 *μ*M Medroxyprogesterone acetate, 10 nM E2, and 50 *μ*M cAMP). Decidual media was changed every 48 h. After 72h of treatment, cells were fixed. Cells were fixed for 15 min with 1/10 volume of freshly prepared formaldehyde solution (11% formaldehyde, 0.1 M NaCl, 1 mM EDTA, and 50 mM HEPES). The fixation was stopped by adding 1/20 volume 2.5 M glycine for 5 min. Fixed DSCs were collected and pelleted at 800 x g for 10 min at 4C. Cell pellets were washed two times with cold PBS-Igepal (0.5% Igepal, 1 mM PMSF). DSCs from six patients were pooled before genomic DNA isolation. GATA2 immunoprecipitation and DNA library generation were performed by Active Motif. ChIP and input DNA were amplified using the Illumina ChIP-Seq DNA Sample Prep Kit. Briefly, DNA ends were polished and 59-phosphorylated using T4 DNA polymerase, Klenow polymerase, and T4 polynucleotide kinase. Addition of 3’-adenine to blunt ends using Klenow fragment (3’-5’ exo minus), Illumina genomic adapters were ligated and the sample was size fractionated to 175–225 bp on a 2% agarose gel. After amplification for 18 cycles with Phusion polymerase, the resulting DNA libraries were tested by RT-qPCR at the same specific genomic regions as the original ChIP DNA to assess quality of the amplification reactions. 36-nt sequence reads were identified by the Sequencing Service (using Illumina’s Hi-Seq). Reads were mapped to the human genome hg19 using the BWA algorithm. Only tags that map uniquely, have no more than 2 mismatches, and that pass quality control filtering are used in the subsequent analysis. The number of unique alignments without duplicate reads was totaled and MACS analysis was performed at a pvalue 1e-7. Raw and processed data are available in SRA and GEO: GSE108409.

### GATA2 knockdown and expression profiling

Three HESC subcultures were transfected with siRNA and treated with decidual media (see above) for 3 days. Total RNA was extracted using the Qiagen RNeasy RNA isolation kit (Qiagen). The RNA from 3 replicates (wells) was pooled for each treatment per cell line. The integrity of all RNA samples was tested with the Bioanalyzer 2100 (Agilent Technologies). The concentration of RNA was quantified on the Nanodrop Spectrophotometer (Nanodrop Technologies). The samples with 260/280 greater than 1.8 were used for microarray hybridization. Microarrays were performed by the Genomic and RNA Profiling Core of Baylor College of Medicine using Affymetrix human genome U133 Plus 2.0 arrays (Affymetrix). Microarray CEL files were analyzed using dChip using the PM-MM model and quantile normalization. Combat was used to normalize differences and for batch correction. Two-side t test and fold changes were used to define differentially expressed genes. Genes with P=0.05 and an absolute fold change 1.4 were considered significant. Raw and processed data are available in SRA and GEO: GSE108409.

### GATA2 ChIP-Seq

ESFs isolated from 6 patients were cultured separately in 150-mm culture dishes with ESF growth media and allowed to reach approximately 90% confluency before being treated with decidual media. Decidual media was changed every 48 hours. After 72 hours of treatment, cells were fixed for 15 minutes with 1/10 volume of freshly prepared formaldehyde solution (11% formaldehyde, 0.1 M NaCl, 1 mM EDTA, and 50 mM HEPES). The fixation was stopped by adding 1/20 volume 2.5 M glycine for 5 minutes. Fixed HESCs were collected and pelleted at 800 g for 10 minutes at 4°C. Cell pellets were washed 2 times with cold PBS, ESFs from 6 patients were pooled before genomic DNA isolation. GATA2 immunoprecipitation and DNA library generation were performed by Active Motif. DNA libraries were sequenced by Illumina’s HiSeq Sequencing Service. 50-nucleotide sequence reads were mapped to the human genome (GRCh Build 37; February 2009) using the Burrows-Wheeler Aligner algorithm with default settings. Alignment information for each read was stored in the Sequence Alignment/Map or Binary version of the Sequence Alignment/Map format. Sequence alignments were extended in silico (using Active Motif software) at their 3 -ends to a length of 150 -250 bp and assigned to 32-nucleotide bins along the genome. The resulting histogram of fragment densities was stored in a binary analysis results file. Raw and processed data are available in SRA and GEO: GSE108409.

### Luciferase assays

A 1000bp region centered on rs2523393 (hg19, chr6:29705520–29706519) was synthesized with either the A or the G allele (Genscript) was cloned into the pGL3-Basic luciferase vector (Promega). A pGL3-Basic plasmid without the 1kb rs2523393 insert was used as a negative control. GATA2 and PGR expression vectors were also obtained from Genscript. Endometrial stromal fibroblasts (ATCC CRL-4003) immortalized with telomerase were maintained in phenol red free DMEM (Gibco) supplemented with 10% charcoal stripped fetal bovine serum (CSFBS; Gibco), 1x ITS (Gibco), 1% sodium pyruvate (Gibco), and 1% L-glutamine (Gibco). Confluent ESFs in 96 well plates in 80*μ*l of Opti-MEM (Gibco) were transfected with 100ng of the luciferase plasmid, 100ng of GATA2 and/or PGR as needed, and 10ng of pRL-null with 0.1μl PLUS reagent (Invitrogen) and 0.25μl of Lipofectamine LTX (Invitrogen) in 20μl Opti-MEM. The cells incubated in the transfection mixture for 6hrs and the media was replaced with the phenol red free maintenance media overnight. Decidualization was then induced by incubating the cells in the decidualization media: DMEM with phenol red (Gibco), 2% CSFBS (Gibco), 1% sodium pyruvate (Gibco), 0.5mM 8-Br-cAMP (Sigma), and 1μM of the progesterone analog medroxyprogesterone acetate (Sigma) for 48hrs. Decidualization controls were incubated in the decidualization control media (phenol red free DMEM (Gibco), 2% CSFBS (Gibco), and 1% sodium pyruvate (Gibco) instead for 48hrs. After decidualization, Dual Luciferase Reporter Assays (Promega) were started by incubating the cells for 15mins in 20μl of 1x passive lysis buffer. Luciferase and renilla activity were then measured using the Glomax multi+ detection system (Promega). Luciferase activity values were standardized by the renilla activity values and background activity values as determined by measuring luminescence from the pGL3-Basic plasmid with no insert. Each luciferase experiment was replicated in at least 4 independent experiments.

### The rs2523393 polymorphism occurs in region that loops to the *HLA-F* promoter

We also took advantage of the fact that rs2523393 was an eQTL for *HLA-F* in EBV transformed lymphocytes (LCLs) to determine if this locus interacts with the *HLA-F* promoter in capture Hi-C (PCHi-C) data generated from LCLs. We used the CHiCP browser ^85^, a web-based tool for the integrative and interactive visualization of promoter capture Hi-C datasets, to identify long range interactions in the high-resolution capture Hi-C dataset of Mifsud et al. from the LCL line GM12878 ^86^.

### The rs2523393 A allele is derived in humans

To reconstruct the evolutionary history of the G/A polymorphism we used a region spanning 50bp upstream and downstream of rs2523393 from hg19 (chr6:32814778–32814878) as a query sequence to BLAT search the chimpanzee (CHIMP2.1.4), gorilla (gorGor3.1), orangutan (PPYG2), gibbon, (Nleu1.0), rhesus monkey (MMUL_1), hamadryas baboon (Pham_1.0), olive baboon (Panu_2.0), vervet monkey (ChlSab1.0), marmoset (C_jacchus3.2.1), Bolivian squirrel monkey (SalBol1.0), tarsier (tarSyr1), mouse lemur (micMur1), and galago (OtoGar3) genomes. For all other non-human species we used the same 101bp region as a query for SRA-BLAST against high-throughput sequencing reads deposited in SRA. The top scoring 100 reads were assembled into contigs using the ‘Map to reference’ option in Geneious v6.1.2 and the human sequence as a reference. Sequences for the Altai Neanderthal, Denisovan, and Ust-Ishim were obtained from the ‘Ancient Genome Browser. The frequency of the G/A allele across the Human Genome Diversity Project (HGDP) populations was obtained from the ‘Geography of Genetic Variants Browser’. We inferred ancestral sequences of the 101bp region using the ancestral sequence reconstruction (ASR) module of Datamonkey^29^ which implements joint, marginal, and sampled reconstruction methods ^30^, the nucleotide alignment of the 101bp, the best fitting nucleotide substitution model (HKY85), a general discrete model of site-to-site rate variation with 3 rate classes, and the phylogeny shown in Fig 6A. All three ASR methods reconstructed the same sequence for the ancestral human sequence at 1.0 support.

## Results

### rs2523393 is a eQTL for *HLA-F*

We have previously shown that the rs2523393 A/G polymorphism is an eQTL for *HLA-F* in mid-secretory phase endometrium^6^. To replicate this observation in an independent cohort and in additional tissues, we tested if rs2523393 was correlated with *HLA-F* expression using GTEx data. Confirming our previous observation, we found that rs2523393 was an eQTL for *HLA-F* in the uterus (P=0.028) as well as 22 other tissues (Fig. 1). We also used GTEx data to identify other genes for which rs2523393 was an eQTL and found that it was an eQTL for 15 other genes, including *HLA-A* and *HLA-G* (**Table S1**). The observation that rs2523393 was an eQTL for *HLA-A* and *HLA-G* in GTEx data prompted us to explore whether it was an eQTL for these genes in the mid-secretory endometrium data from our original study^6^. Indeed we found that rs2523393 was also an eQTL for *HLA-G* (P=0.041), but not *HLA-A* in mid-secretory endometrium.

**Fig 1.**
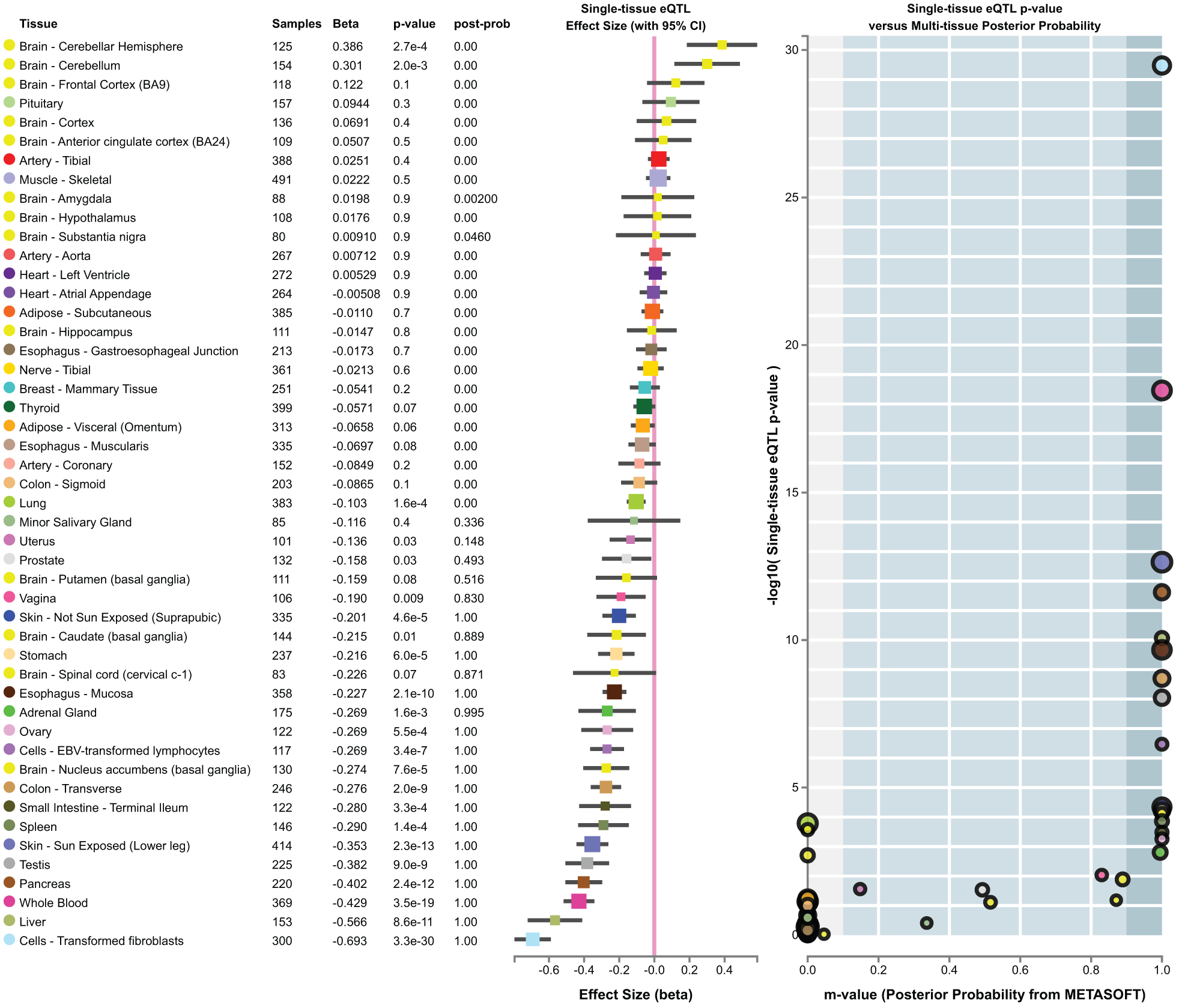
GTEx multi-tissue eQTL plot for rs2523393. Left, Tissue type, sample size, beta, p-value, and posterior probability that rs2523393 in each tissue tested is also an eQTL across other tissues. Center, single-tissue effect size and 95% confidence interval. Right, single-tissue eQTL p-value versus multi-tissue posterior probability.

### The rs2523393 G/A polymorphism occurs within a GATA2 motif

To infer the functional consequences of the G/A polymorphism we used DeepSea ^31^, a deep learning-based algorithm that infers the effects of single nucleotide substitutions on chromatin features such as transcription factors binding, DNase I sensitivity, and histone marks. DeepSea predicted the G allele would have a negative effect on the binding of GATA2 (log2 fold change effect: -2.07, E-value: 0.035) and GATA3 (log2 fold change effect: -1.00, E-value: 0.003) (Fig 2A). We next used JASPAR transcription factor binding site (TFBS) motifs ^32^ to identify putative TFBSs in a 36bp window upstream and downstream of rs2523393. We identified a single TFBS in this window, a GATA motif (matrix ID: MA0036.3, score: 8.70) and found that the G/A polymorphism occurs at an invariant A site in the motif (GATAA). Similar to the DeepSea results, substituting the reference A allele with the alternative G allele is predicted to abolish GATA2 binding (Fig 2B/C). These results suggest that the G/A polymorphism alters binding of GATA2, a transcription factor that plays an essential role in mediating the transcriptional response to progesterone in decidual stromal cells (DSCs)^33–35^.

**Fig 2.**
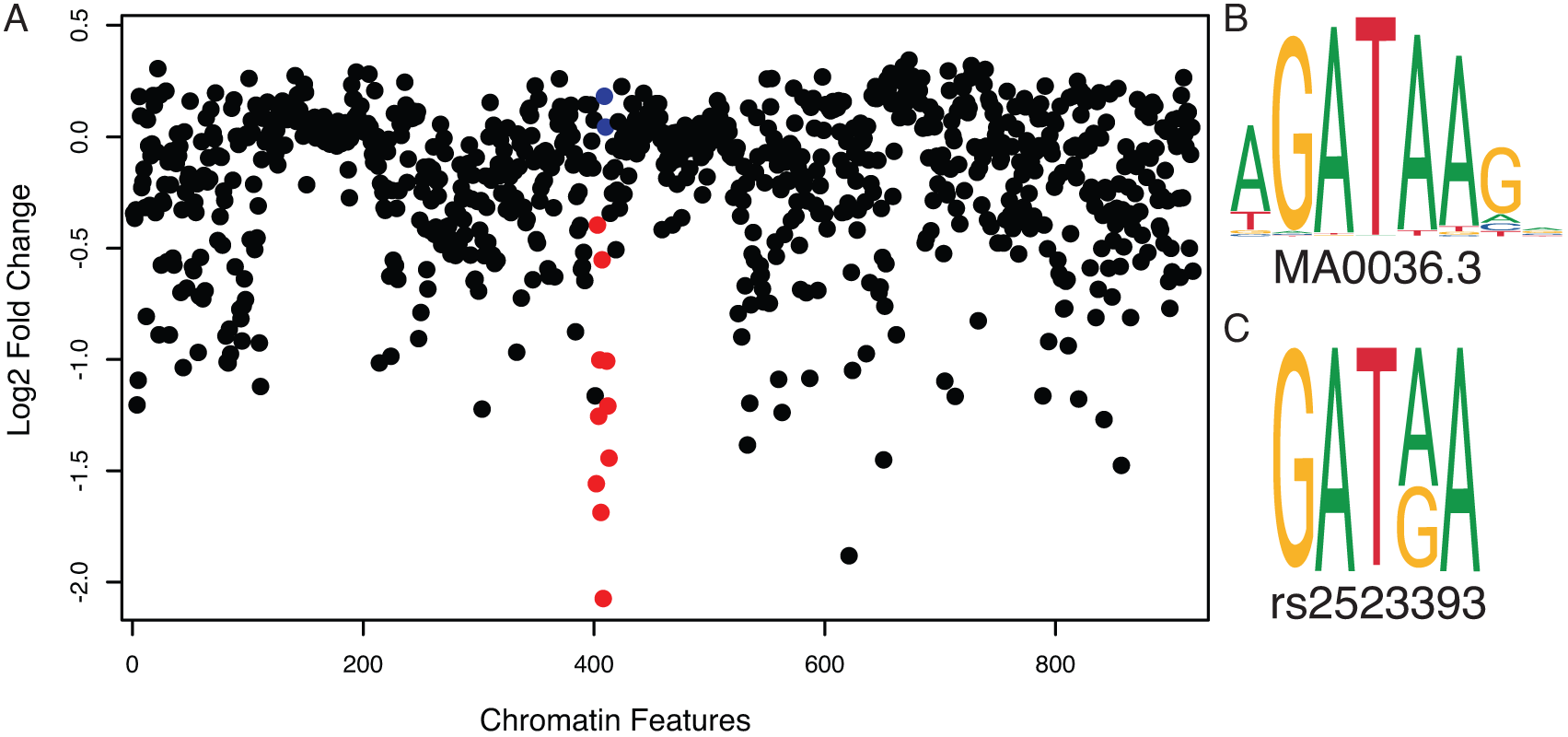
The rs2523393 G allele is predicted to abolish a GATA2 binding site. (**A**) DeepSea plot showing predicted effects of the G allele on chromatin features. GATA1 binding, blue. GATA2 and GATA3 binding, red. (**B**) JASPAR GATA2 motif. (**C**) GATA2 motif at rs2523393.

### *HLA-F* is up-regulated by progesterone and GATA2 in decidual stromal cells

Previous studies have shown that HLA-F is expressed in placental villi, extra-villus trophoblasts (EVT), and DSCs ^9–11,36^. To determine if the expression of *HLA-F* varies in the endometrium across the menstrual cycle, we used previously generated microarray expression data ^21^. We found that *HLA-F* expression reached peak expression in the late secretory phase of the menstrual cycle, and was significantly differentially expressed throughout the menstrual cycle (Fig 3A). To test if *HLA-F* is up-regulated by progesterone, we examined its expression in RNA-Seq data from ESFs treated with control media or differentiated with cAMP/progesterone into DSCs treated with either a PGR specific siRNA or a scrambled control siRNA ^26,27,37^. We found that *HLA-F* was up-regulated 2.11-fold (P=3.59×10^−5^; Fig 3B) in DSCs and that knockdown of PGR down-regulated *HLA-F* 1.67-fold (P=6.15×10^−8^; Fig 3B). Next we used an siRNA to knockdown the expression of GATA2 in DSCs and assayed global gene expression using an Agilent Human Gene Expression 8x60K microarray. Consentient with our previous results, we observed a 2.20-fold down-regulation of HLA-F (P=0.02) upon GATA2 knockdown (Fig 3C). We also observed that HLA-F expression was lower in the endometria of women with implantation failure (1.38-fold, *P*=1.54×10^−3^) or recurrent spontaneous abortion (1.35-fold, 0.011) compared to fertile controls (Fig 3D). Thus we conclude that *HLA-F* expression increases as the menstrual cycle progresses, is up-regulated as by progesterone, PGR, and GATA2 during the differentiation of ESFs into DSCs, and dysregulation is associated with implantation failure and recurrent spontaneous abortion.

**Fig 3.**
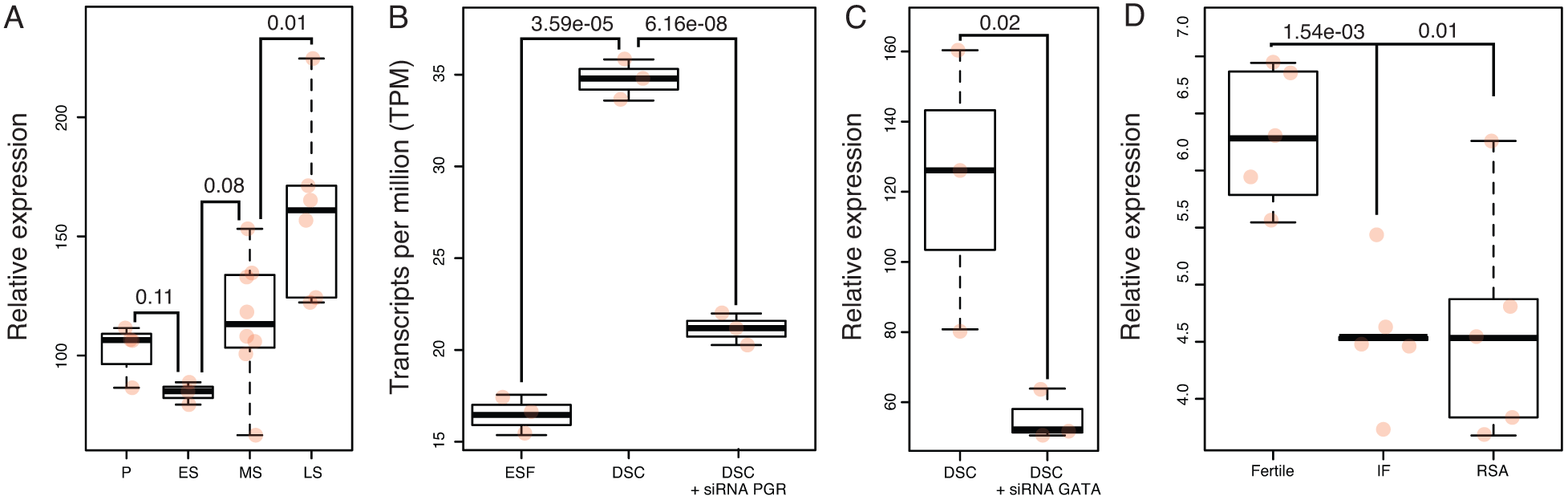
HLA-F is regulated by upregualted progesterone and GATA2 during the menstrual cycle and in human decidual stromal cells. (A) Relative expression of HLA-F throughout the menstrual cycle. P, proliferative phase. ES, early secretory phase. MS, mid-secretory phase. LS, late secretory phase. N=3–8, T-test. (B) Expression level of HLA-F in endometrial stromal fibroblasts (ESF), decidual stromal cells (DSCs), and DSCs treated with an siRNA to knockdown PGR expression. (C) Relative expression level of HLA-F in DSCs and DSCs treated with an siRNA to knockdown GATA2 expression. N=3, T-test. (D) Relative expression level of HLA-F in the endometrial fertile women, and women with implantation failure (IF) or recurrent spontaneous abortion (RSA) N=5, T-test.

### The rs2523393 A allele creates a new enhancer that loops to the *HLA-F*

Our observations that rs2523393 is an eQTL for *HLA-F* and that progesterone and GATA2 up-regulate *HLA-F* in DSCs suggests rs2523393 may be located within or linked to a progesterone responsive enhancer. To identify such a regulatory element we generated new GATA2 ChIP-Seq data from DSCs and used previously published ChIP-Seq data from DSCs for the transcription factors PGR ^27,38,39^ FOXO1^38^, FOSL2^38^, and NR2F2 (COUP-TFII) ^40^, H3K27ac which marks active enhancers^27^, H3K4me3 which marks active promoters ^27^, and DNaseI-Seq and FAIRE-Seq to identify regions of open chromatin ^27^. We also took advantage of the fact that rs2523393 was an eQTL for *HLA-F* in multiple tissues, including EBV transformed lymphocytes (LCLs), to identify additional transcription factor binding sites using ChIP-Seq data generated by ENCODE and promoter capture Hi-C (PCHi-C) data generated from LCLs. We found that rs2523393 was located in a region of open chromatin and within a GATA2 ChIP-Seq peak in DSCs, 170bp upstream of a FOSL2 ChIP-Seq peak in DSCs, and nearby a cluster of transcription factor binding sites in ENCODE data including a binding site for GATA2 (Fig 4). Finally, we found that this region loops to the *HLA-F* promoter in LCLs located ~14.5kb upstream of rs2523393, suggesting this region is an enhancer for HLA-F.

**Fig 4.**
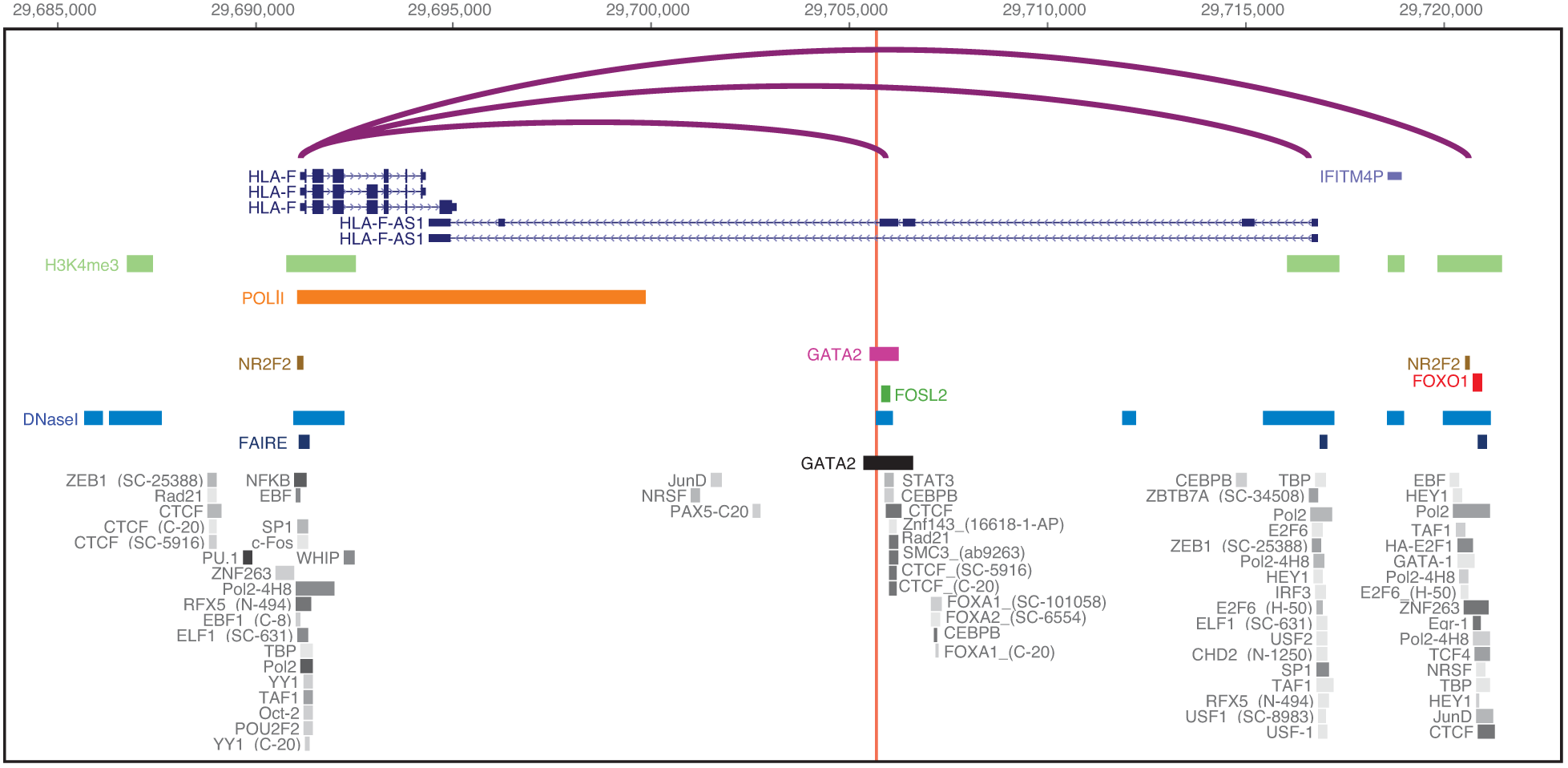
The rs2523393 G/A polymorphism is located in a distal enhancer of HLA-F. Location of rs2523393 polymorphism (vertical red line) with respect to histone modifications that characterize promoters (H3K4me4 ChIP-Seq), enhancers (H3K27ac ChIP-Seq), open chromatin (DNaseI-Seq, FAIRE-Seq), as well as GATA2 and NR2F2 ChIP-Seq binding sites in human DSCs. ENCODE transcription factor binding sites are shown in grey and intrachromosomal loop (PCHiC) data from LCLs is shown in purple.

To test if this locus has regulatory potential we synthesized a 1000bp region spanning 500bp upstream and downstream of rs2523393 (Fig 4) with either the reference A allele or the alternate G allele and cloned them into the pGL3-Basic[minP] luciferase reporter vector, which lacks an enhancer but encodes a minimal promoter. Next we transiently transfected either the pGL3-Basic[minP]-rs2523393A or pGL3-Basic[minP]-rs2523393G luciferase reporter along with the pRL-null internal control vector into DSCs and quantified luciferase and renilla activity using a dual luciferase assay. We found that luciferase activity was significantly lower in DSCs transfected either pGL3-Basic[minP]-rs2523393A (Wilcox test *P*=3.80×10^−4^) or pGL3-Basic[minP]-rs2523393G (Wilcox test *P*=3.82×10^−5^) compared to empty vector pGL3Basic[minP] controls (Fig 5). Next, we co-transfected pGL3-Basic[minP]-rs2523393A or pGL3-Basic[minP]-rs2523393G along either a GATA2 expression vector or a PGR expression vector. While co-transfection of neither the GATA2 expression vector nor the PGR expression vector enhanced luciferase expression above empty vector controls, there were significant differences between alleles (Fig 5). Finally we co-transfected pGL3-Basic[minP]-rs2523393A or pGL3-Basic[minP]-rs2523393G with both GATA2 and PGR expression vectors. Consistent with cooperative interaction between GATA2 and PGR, we observed a significant increase in luciferase expression from both the pGL3-Basic[minP]-rs2523393A (Wilcox test *P*=3.00×10^−10^) and pGL3-Basic[minP]-rs2523393G (Wilcox test *P*=3.61×10^−7^) vectors compared to empty vector controls. We also observed that the pGL3-Basic[minP]-rs2523393A vector drove significantly higher luciferase expression than the pGL3-Basic[minP]-rs2523393G vector (Wilcox test *P*=3.65×10^−3^). Thus we conclude that rs2523393 is located within a GATA2 binding site in a progesterone responsive enhancer, and that the G/A polymorphism affects enhancer function.

**Fig 5.**
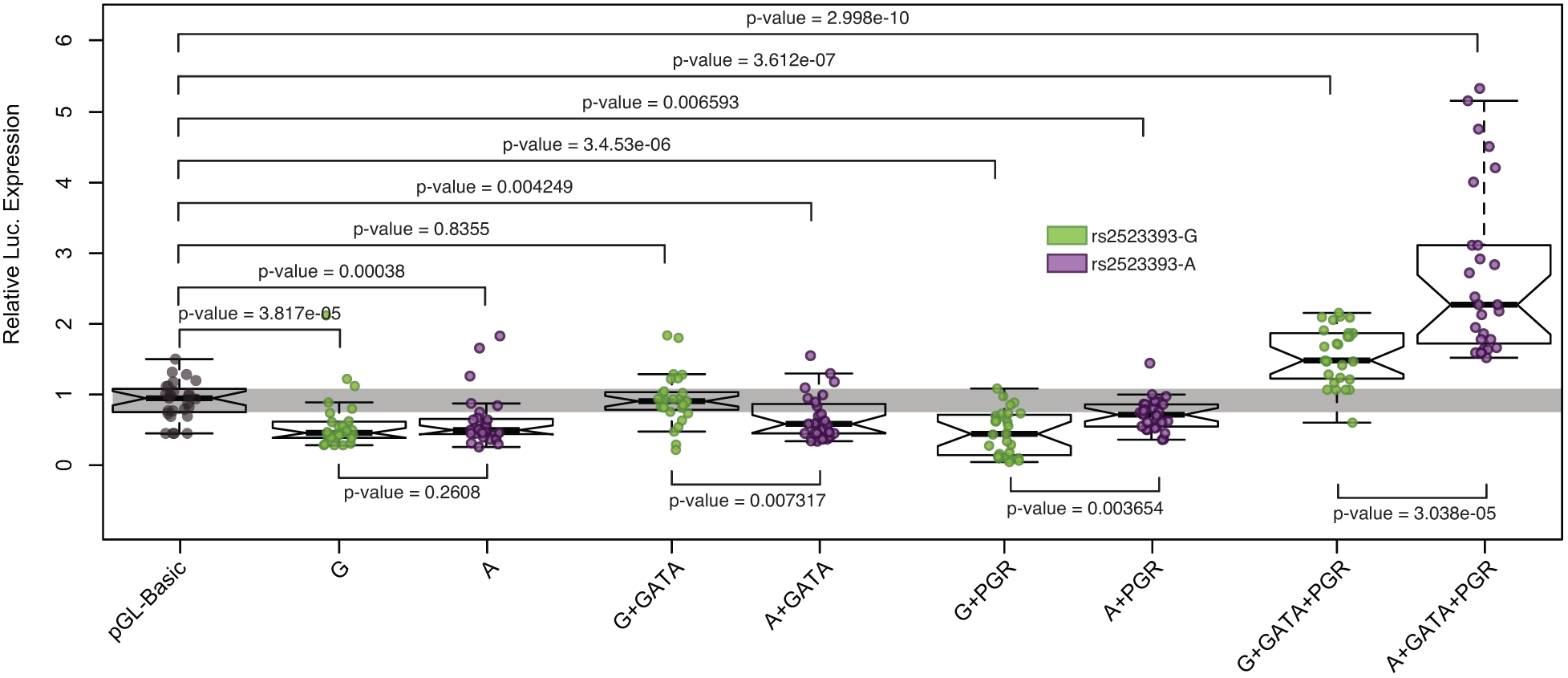
The rs2523393 G/A polymorphism alters the function of a progesterone response enhancer. Luciferase assay results testing the regulatory potential of the G (pGL3Basic-rs2523393G) and A (pGL3Basic-rs2523393A) alleles in DSCs treated with cAMP/MPA for 48 hours. Data are shown as luciferase activity from the pGL3Basic-rs2523393G or pGL3Basic-rs2523393A reporter relative to renilla activity (pRL-null and empty vector (pGL3Basic) controls. +GATA, Relative luciferase activity from the pGL3Basic-rs2523393G or pGL3Basic-rs2523393A in DSCs co-transfected with GATA. +PGR, Relative luciferase activity from the pGL3Basic-rs2523393G or pGL3Basic-rs2523393A in DSCs co-transfected with PGR. +GATA & PGR, Relative luciferase activity from the pGL3Basic-rs2523393G or pGL3Basic-rs2523393A in DSCs co-transfected with GATA and PGR. n=14, p=Wilcox test.

### The rs2523393 A allele is derived in humans

To reconstruct the evolutionary history of the G/A polymorphism we identified a region spanning 50bp upstream and downstream of rs2523393 from 37 primates, including species from each of the major primate lineages, multiple sub-species of African apes (Homininae), as well as modern and archaic (Altai Neanderthal and Denisova) humans. Next we used maximum likelihood methods to reconstruct ancestral sequences for this 101bp region. We found that the G allele was ancestral in primates and that the A variant was only found in Neanderthal, Densiovan, and modern human populations (Fig 6A). Next we examined the frequency of these alleles across the Human Genome Diversity Project (HGDP) populations and found that the derived and ancestral alleles were segregating at intermediate to high frequencies in nearly all human populations (Fig 6B). These results indicate that the derived A variant at rs2523393, which creates a new GATA2 binding site in an enhancer of HLA-F expression, arose before the population split between archaic and modern humans between 550,000 and 765,000 years ago^41^.

**Fig 6.**
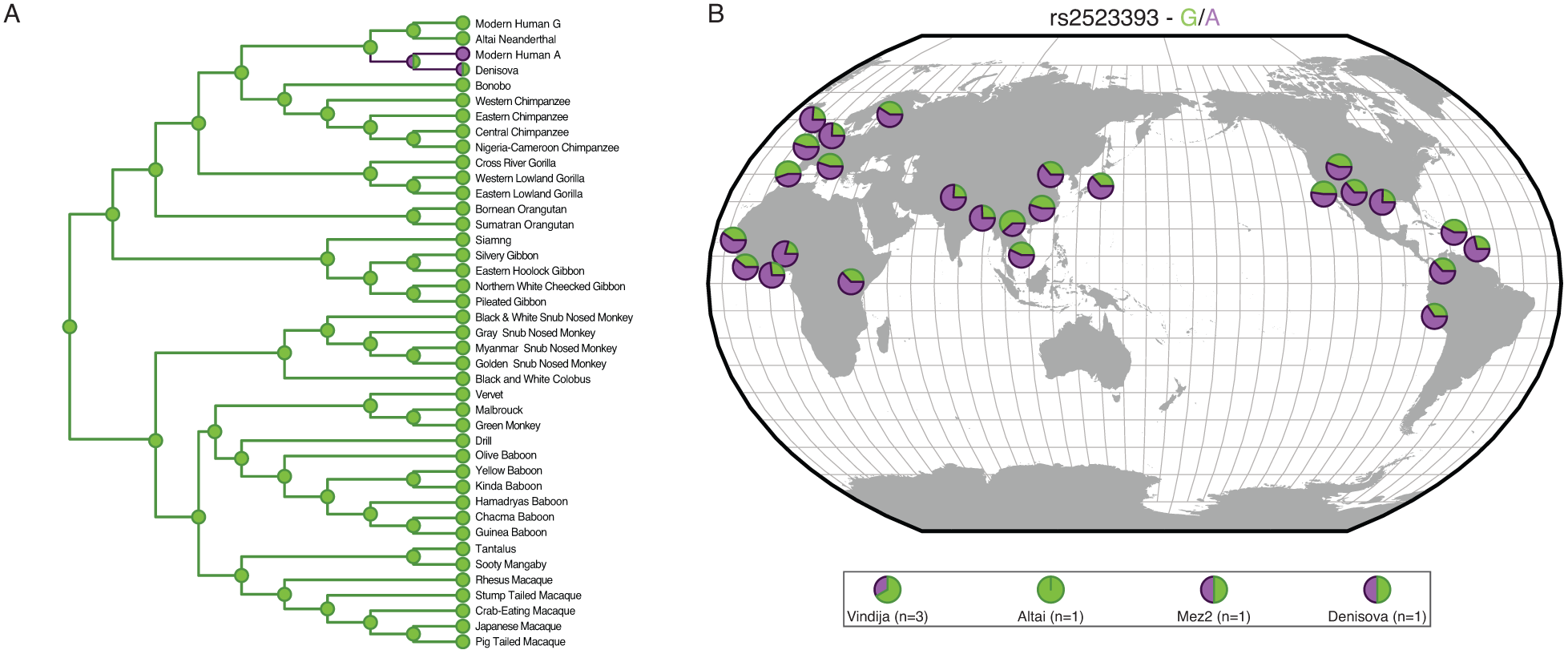
Evolutionary history G/A polymorphism at rs2523393. (**A**) rs2523393 genotype across primates and ancestral reconstructions. Genotype in extant species is shown as circles next to species names at terminal branches. Genotype at internal nodes based on ancestral reconstruction is shown as circles. Purple, A. Green, G. (**B**) Distribution of the G and A alleles of rs2523393 across HGDP populations.

### The rs2523393 G/A polymorphism is associated with disease

Our observation that the rs2523393 G/A polymorphism is relatively old and segregating at intermediate to high frequencies implies the polymorphism is being maintained by balancing selection, frequency dependent selection, or antagonistic pleiotropy ^42^. Intriguingly, a previous GWAS found the rs2523393 A allele was associated with multiple sclerosis (MIM: 126200) with an odds ratio of 1.28 [1.18–1.39] (P=1.04×10^−17^)^43^ suggesting antagonistic pleiotropy maintains both alleles. To explore if other disease were associated with rs2523393, we took advantage of the UK Biobank GWAS results of ~2,000 phenotypes (Fig 7). We found that the rs2523393 G allele was most significantly associated with malabsorption/coeliac disease (P=8.8×10^−24^, beta =-0.0016), multiple sclerosis (P=2.7×10^−10^, beta =-0.0009), and hyperthyroidism/thyrotoxicosis (P=1.2×10^−8^, beta =-0.0012).

**Fig 7.**
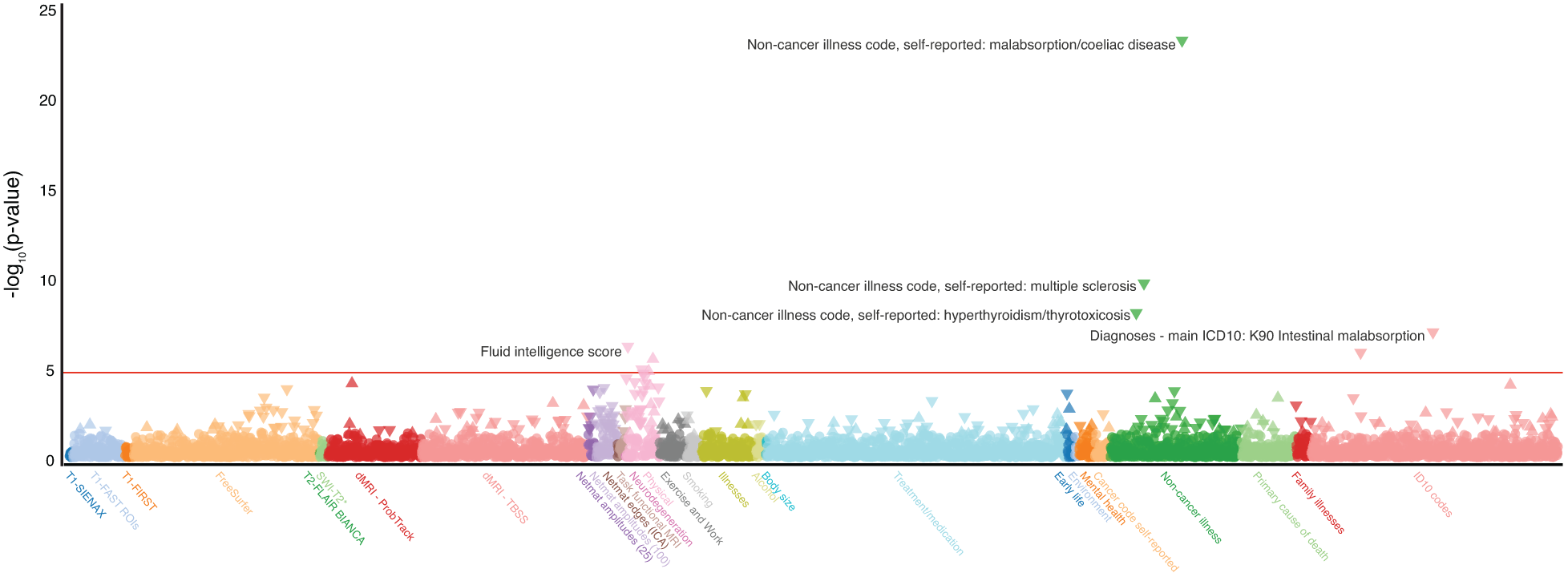
PheWAS plot of phenotypes associated with rs2523393 in the UK Biobank. Associations are shown for 3870 ICD10 codes, associations with P<0.05 are shown are triangles and those with P≥0.05 are shown as circles. The most statistical significant phenotype associations are labled.

## Discussion

The mechanisms that underlie maternal tolerance of the antigenically distinct fetus are complex ^44^ and have been the subject of intense study since Medawar proposed tolerance was achieved by maternal immunosuppression and immaturity of fetal antigens ^45^. It is now clear that rather than being suppressed at the maternal-fetal interface, the immune system plays an active role in establishing a permissive environment for implantation, placentation, and gestation. Among the diverse immune cells that contribute to the successful establishment and maintenance of pregnancy are maternal regulatory T cells (Tregs) and γδ T cells ^46–48^, uterine natural killer cells ^49,50^, uterine dendritic cells ^51^, uterine macrophage ^52^, as well as DSCs themselves ^53–55^. DSCs, for example, recruit uterine natural killer cells to the endometrium via IL-15 expression ^56–64^, stimulate the migration of macrophage into the endometrium via CSF1 expression ^56–58,65,66^, and suppress cytokine secretion by allogenic CD4+ T cells via PD-1 ligand interactions^54^, which are all essential for establishing maternal-fetal immunotolerance. These data indicate that DSCs directly regulate local immune responses through multiple mechanisms, potentially also through HLA-F.

Major histocompatibility complex (MHC) genes play an important role in the rejection of non-self tissues, but also contribute to maternal tolerance of the fetus ^1–3,5,67–69,4,70–81^. While the precise functions of HLA-F are unclear, our observations that its expression level in the endometrium during the window of implantation is associated with fecundability^6^, and that it is up-regulated by progesterone implicates it in the establishment of pregnancy. Intriguingly HLA-F binds LIR and KIR natural killer cell receptors^13–17^, suggesting HLA-F expressed by DSCs directly signal to uterine natural killer (uNK) cells which are essential for the establishment of maternal immunotolerance and spiral artery remodeling. Consistent with this hypothesis HLA-F expression level is positively correlated with uNK abundance in mid-luteal endometria and is predictive of achieving pregnancy^6,18^.

Our observation that the G/A polymorphism is shared between modern humans, Neanderthals, and Denisovans indicates it is relatively ancient and may be maintained by balancing selection, frequency dependent selection, or antagonistic pleiotropy. We found that while the derived A allele creates a new GATA2 binding site and augments the function of a progesterone responsive enhancer of HLA-F, a previous GWAS found it was also associated with multiple sclerosis ^43^ and the G polymorphism is also associated with malabsorption/coeliac disease, multiple sclerosis, and hyperthyroidism/thyrotoxicosis in the UK Biobank. These data are consistent with maintenance through antagonistic pleiotropy. We have previously shown that another fecundability-associated variant, which switches a repressor into an enhancer of endometrial *TAP2* expression, was also shared between modern humans, Neanderthals, and Denisovans^82^. Remarkably, while the ancestral T allele in the *TAP2 cis*-regulatory element was associated with shorter time to pregnancy and ulcerative colitis ^83^, the derived C allele was associated with longer time to pregnancy and Crohn’s disease (MIM: 266600) ^84^ suggesting theses alleles are also maintained by antagonistic pleiotropy. Collectively these data suggest there is a reproduction-health tradeoff in human evolution.

## Acknowledgments

This work was funded by a Burroughs Wellcome Preterm Birth Initiative grant (1013760), an NIH National Institute of General Medical Sciences Graduate Training Grant (T32GM007197), and by the March of Dimes Transdisciplinary Center (TDC) at UChicago-Northwestern-Duke.

## Web Resources

GTEx database (http://www.qtexportal.orq/home/eqtls/bvSnp), GEO2R analysis package (http://www.ncbi.nlm.nih.gov/qeo/qeo2r/), Geography of Genetic Variants Browser (http://popgen.uchicago.edu/ggv/), Datamonkey web-sever (http://www.datamonkev.org), Online Mendelian Inheritance in Man (http://www.omim.org), Oxford Brain Imaging Genetics (BIG) Server (http://big.stats.ox.ac.uk), CHiCP, a web-based tool for the integrative and interactive visualization of promoter capture Hi-C datasets (https://www.chicp.org).

